# Rab10-positive tubular structures represent a novel endocytic pathway that diverges from canonical macropinocytosis in RAW264 macrophages

**DOI:** 10.1101/2021.01.03.425161

**Authors:** Katsuhisa Kawai, Arata Nishigaki, Seiji Moriya, Youhei Egami, Nobukazu Araki

## Abstract

Using the optogenetic photo-manipulation of photoactivatable (PA)-Rac1, remarkable cell surface ruffling and the formation of a macropinocytic cup (premacropinosome) could be induced in the region of RAW264 macrophages irradiated with blue light due to the activation of PA-Rac1. However, the completion of macropinosome formation did not occur until Rac1 was deactivated by the removal of the light stimulus. Following PA-Rac1 deactivation, some premacropinosomes closed into intracellular macropinosomes, whereas many others transformed into long Rab10-positive tubules without forming typical macropinosomes. These Rab10-positive tubules moved centripetally towards the Golgi region along microtubules. Surprisingly, these Rab10-positive tubules did not contain any endosome/lysosome compartment markers, such as Rab5, Rab7, or LAMP1, suggesting that the Rab10-positive tubules were not part of the degradation pathway for lysosomes. These Rab10-positive tubules were distinct from recycling endosomal compartments, which are labeled with Rab4, Rab11, or SNX1. These findings suggested that these Rab10-positive tubules belonged to a novel, non-degradative, endocytic pathway. The formation of Rab10-positive tubules from premacropinosomes was also observed in control and phorbol myristate acetate (PMA)-stimulated macrophages, although their frequencies were low. Interestingly, the formation of Rab10-positive premacropinosomes and tubules was not inhibited by phosphoinositide 3-kinase (PI3K) inhibitors, while the classical macropinosome formation requires PI3K activity. Thus, this study provides evidence to support the existence of Rab10-positive tubules as a novel, non-degradative, endocytic pathway that diverges from canonical macropinocytosis.

## Introduction

Macropinocytosis is a form of clathrin-independent, actin-dependent endocytosis that mediates the non-selective uptake of extracellular fluids and solutes into 0.2 – several μm diameter endocytic vesicles called macropinosomes. Newly-formed macropinosomes undergo a maturation process, during which they fuse with other endocytic compartments and eventually merged with lysosomes (1,2). In macrophages and dendritic cells, this endocytic pathway is involved in immune surveillance and antigen presentation (3). In tumor cells, macropinocytosis serves as an amino acid supply route to support active cell proliferation (4). In neuronal cells, macropinocytosis has been proposed to be involved in the uptake and propagation of pathogenic protein aggregates associated with neurodegenerative diseases such as, Alzheimer’s disease and Parkinson’s disease (5). In addition, many pathogenic bacteria and viruses exploit macropinocytosis pathways to gain entry into host cells during infection (6). Therefore, a better understanding of the molecular mechanisms underlying macropinocytosis and its related pathways has implications for cell biology and the establishment of therapeutic strategies to combat various diseases, including cancer and viral infections.

Phosphoinositide metabolism and small GTPases of the Ras superfamily, including the Rho and Rab families, coordinately regulate macropinosome formation and maturation (7,8). Macropinocytosis is initiated by the plasma membrane ruffle formation driven by the actin polymerization and reorganization which is upregulated by Rac1, a member of the Rho family GTPases. Using an optogenetic technology in which RAW macrophages express photoactivatable Rac1 (PA-Rac1), we have found that Rac1 activation is sufficient to induce plasma membrane ruffling and circular ruffle/macropinocytic cup formation; however, persistent Rac1 activation stalls at the macropinocytic cup stage without progressing to the complete formation of macropinosomes. Rac1 deactivation following temporal activation enables some macropinocytic cups to form macropinosomes (9). By repeating the activation and deactivation of PA-Rac1 at a certain interval, macropinocytosis can be efficiently induced. However, many macropinocytic cups collapse prior to the completion of macropinosome formation. It is also known that a considerable number of circular ruffles or macropinocytic cups in macrophages under normal conditions disappear without resulting in intracellular macropinosome formation (1). Nonetheless, no attention has been paid to the phenomenon of cup collapse so far. Therefore, in this study, we focused our attention on the differences between collapsing macropinocytic cups and typical cups that form macropinosomes.

The Rab family GTPases comprise more than 60 members in mammals and are key regulators of membrane trafficking (10,11). One protein, Rab10, has been implicated in a variety of membrane trafficking pathways (12–16). More recently, Rab10 was identified as a novel protein regulator of tubular recycling endosome formation through interaction with the kinesin motor protein KIF13 (17). Furthermore, it was reported that Rab10 phosphorylation by leucine-rich repeat kinase 2 (LRRK2) has been associated with the regulation of macropinosome early maturation (18).

In this study, while studying RAW264 macrophages expressing a fluorescent protein-tagged Rab10 and PA-Rac1, we unexpectedly discovered that macropinocytic cup or pocket-like structures (premacropinosomes) that were intensively associated with Rab10 collapsed and disappeared by transforming into Rab10-positive tubules, whereas those premacropinosomes that were only faintly or briefly associated with Rab10 became intracellular macropinosomes. The Rab10-positive tubules that originated from premacropinosomes were distinct from known endocytic compartments having Rab4, Rab5, Rab7, Rab11, SNX1, or lysosomal associated membrane protein 1 (LAMP1). Here, we provide evidence to support the existence of a new endocytic pathway that diverges from canonical macropinocytosis.

## Results

### Reversible optogenetic control of PA-Rac1 activity induces Rab10-positive tubule formation

Through the optogenetic control of PA-Rac1 activity in RAW264 cells, we were able to manipulate the process of macropinocytosis (9). The activation of PA-Rac1 by local irradiation with blue light induced cell surface ruffling. Continuous blue-light irradiation applied to the same area results in the formation of numerous bubble-like structures underneath the ruffles. Our previous study revealed that these bubble-like structures are unclosed macropinocytic cups or pockets, which we refer to as premacropinosomes. After turning off the blue-light irradiation, some premacropinosomes were closed into intracellular macropinosomes, but many other premacropinosomes (~70%) disappear without forming macropinosomes (Figure 1A, Supplementary Movie 1), as we have previously reported (9).

**Figure 1.**
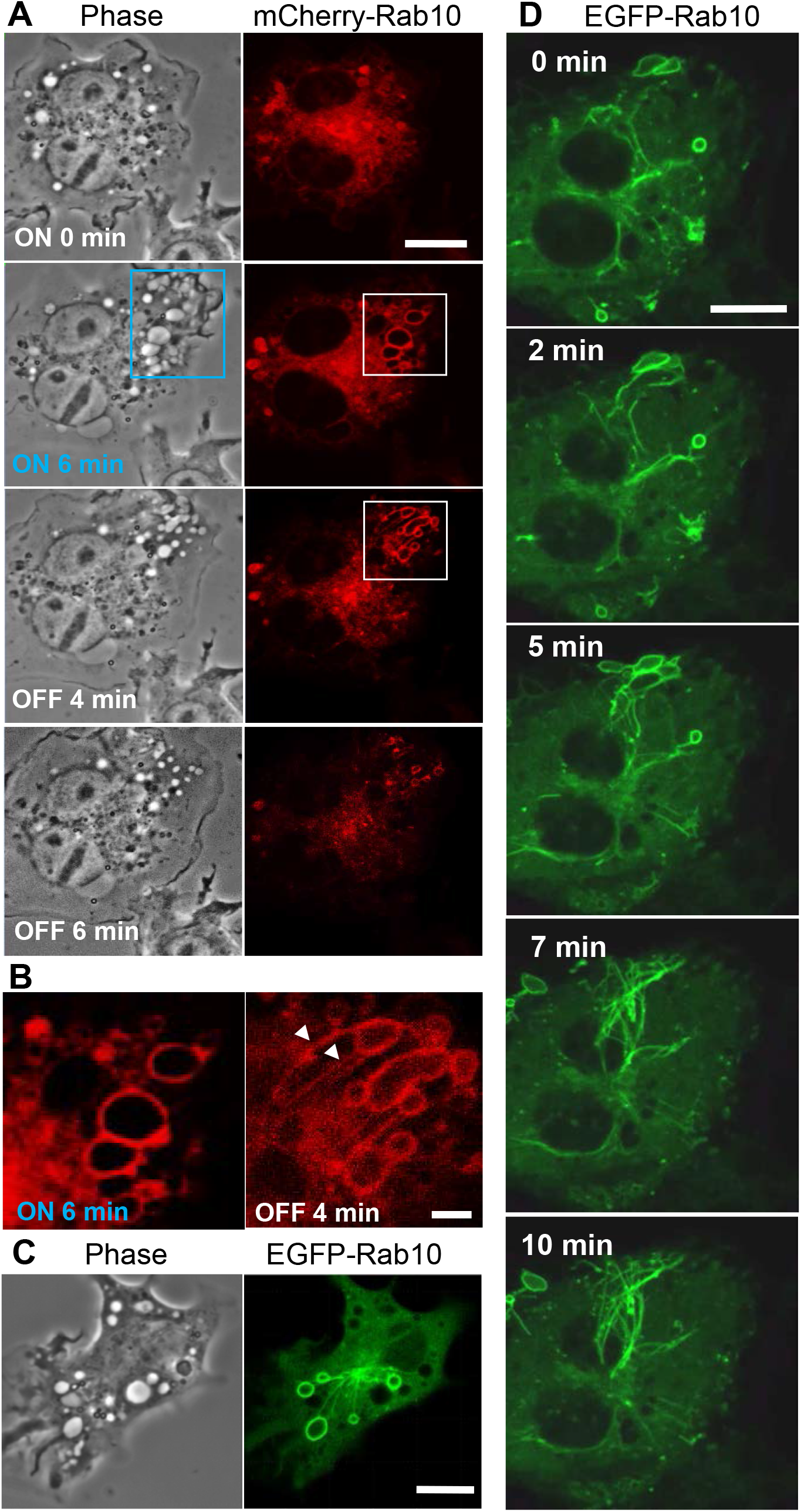
Rab10 dynamics in live RAW264 cells under the optogenetic control of PA-Rac1 activity. **(A)** The blue boxed area of a cell expressing CFP-PA-Rac1 was irradiated with a blue laser to photoactivate PA-Rac1. Phase-contrast (left) and mCherry-Rab10 (right) images were acquired at the indicated times after PA-Rac1 activation (ON) and deactivation (OFF). Scale bar =10 μm. (B) Enlarged images of the white boxed area in **A**. Following local PA-Rac1 activation, Rab10-positive premacropinosomes were formed in the area. After PA-Rac1 OFF, a few Rab10-positive tubules extended from a premacropinosome (arrowheads). Scale bar = 2 μm. **(C)** Representative phase-contrast and EGFP-Rab10 fluorescence images of RAW264 cells during PA-Rac1 ON-OFF cycles. (D) Confocal time-lapse microscopy of EGFP-Rab10 in RAW264 cells during PA-Rac1 ON-OFF cycles. Selected frames from the time-lapse movie are presented. Elapsed times are shown in the frame. Scale bar = 10 μm. The corresponding movie is available in the Supplementary Materials Movie 2.

To clarify the involvement of Rab10 in this process, enhanced green fluorescent protein (EGFP) -or mCherry-tagged Rab10 was co-expressed together with enhanced cyan fluorescent protein (ECFP)-tagged PA-Rac1 and observed in live RAW264 cells during the optogenetic control of PA-Rac1 activity. We found that most PA-Rac1induced premacropinosomes were intensively positive for Rab10. Following the deactivation of PA-Rac1, these Rab10-positive premacropinosomes transformed into tubular structures. Under phase-contrast microscopy, premacropinosomes appeared to collapse and return to the cell surface without forming phase-bright macropinosomes; however, they actually transformed into tubular structures. Curiously, only Rab10-negative premacropinosomes, ~30 % of all premacropinosomes, became phase-bright macropinosomes (Figures 1A, B, Supplementary Figures S1).

When we observed EGFP-Rab10 dynamics in RAW264 cells co-expressing with mCherry-PA-Rac1, which was temporally activated by 488 nm excitation for EGFP-Rab10 observation with 15-sec intervals, we found that the Rab10-positive tubules frequently form from the Rab10-positive premacropinosomes (Figures 1C, D). By live-cell imaging of RAW264 cells co-expressing EGFP-Rab10 with PA-Rac1, it was apparently shown that a few Rab10-tubules extended from one peripheral premacropinosome toward the cell center perinuclear region. The body of the premacropinosome gradually shrunk and disappeared without moving into the cell center (Figure 1D, Supplementary Movie 2). We were able to constantly induce the formation of Rab10-positive tubules from multiple premacropinosomes over a 30-min observation period through repeating PA-Rac1 activation (ON) and deactivation (OFF) cycles with the 488 nm-excitation light for EGFP observation.

### The Rab10-positive tubular endocytic pathway exists in cells under near-physiological conditions

Rab10-positive tubule may reflect an artificial structure that is an artifact of PA-Rac1 photo-manipulation. To rule out this possibility, we examined whether Rab10-positive tubules could be observed in RAW264 cells under near-physiological conditions. In RAW264 cells stimulated with 10 nM phorbol myristate acetate (PMA), membrane ruffling and macropinosome formation were enhanced. During the normal process of macropinosome formation, Rab10 transiently localized to nascent macropinosomes. The Rab10 localization to macropinosomal membranes was much sparser than that on premacropinosomes formed by the photo-manipulation of PA-Rac1 ON-OFF cycling. During a long and careful observation, we observed that Rab10-positive premacropinosomes and tubules were occasionally formed in PMA-stimulated cells (Figures 2, Supplementary Movie 3). Counting the number of Rab10-positive premacropinosomes and Rab10-negative macropinosomes formed showed that the percentage of Rab10-positive premacropinosomes across all macropinosomal structures was ~23 %, which is a much lower proportion than was observed in RAW cells during the photo-manipulation of PA-Rac1 ON-OFF cycling (~68%) (Supplementary Figure S1C). Similarly, the formation of Rab10-positive premacropinosomes was also observed in control RAW264 cells, although the frequency of this evet is lower than in PMA-stimulated cells (Supplementary Figure S1C). Thus, Rab10-positive premacropinosomes and tubules exist in cells under near-physiological conditions, although they were likely easily overlooked due to their shorter lifetimes and lower frequency of appearance. They disappeared within a few minutes by transforming to tubular profiles.

**Figure 2.**
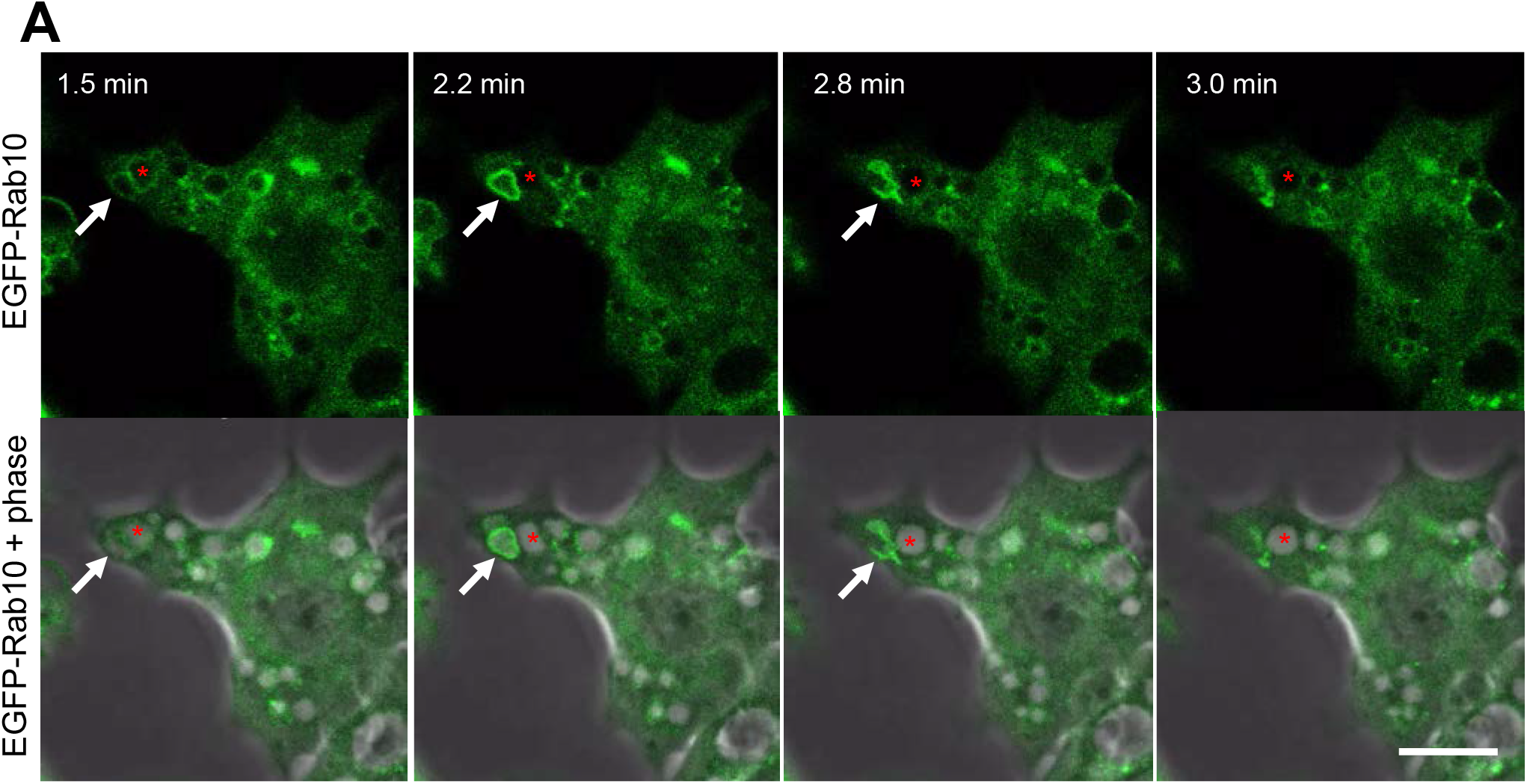
Short-lived Rab10-positive premacropinosomal structures exist in RAW264 cells under near-physiological conditions. Confocal live-cell imaging of RAW 264 cells expressing EGFP-Rab10 stimulated with 10 nM phorbol myristate acetate (PMA). Elapsed times after the addition of PMA are shown in the frame. Arrows indicate a Rab10-positive premacropinosome which disappears within a few minutes. Asterisks indicate a Rab10-negative macropinosomal structure that remains as a phase-bright macropinosome. Scale bar = 10 μm.

Compared with near-physiological conditions, the frequency of Rab10-positive premacropinosome formation significantly increases under photo-manipulation conditions, therefore, we attempted to characterize these Rab10-positive premacropinosomes and tubules mainly under the condition of repeating PA-Rac1 ON-OFF cycles.

### Rab10-positive tubules originate from unclosed macropinocytic cups

To examine whether Rab10-positive macropinosomal structures are closed macropinosomes or unclosed premacropinosomes, the lipophilic dye FM4-64 was added to the cells immediately after confirming the formation of EGFP-Rab10-positive tubules by the photo-manipulation of PA-Rac1 ON-OFF cycling. The cell surface membrane and unclosed premacropinosomes were labeled with FM4-64, whereas phase-bright intracellular macropinosomes are not labeled. Examining EGFP-Rab10 localization, we found that all Rab10-positive premacropinosomes were labeled with FM4-64. The Rab 10-positive tubules that elongated from premacropinosomes were labeled with the FM4-64 dye, which indicated that Rab10-positive tubules elongated directly from the premacropinosomes, which were continuous with the cell surface plasma membrane. (Figure 3).

**Figure 3.**
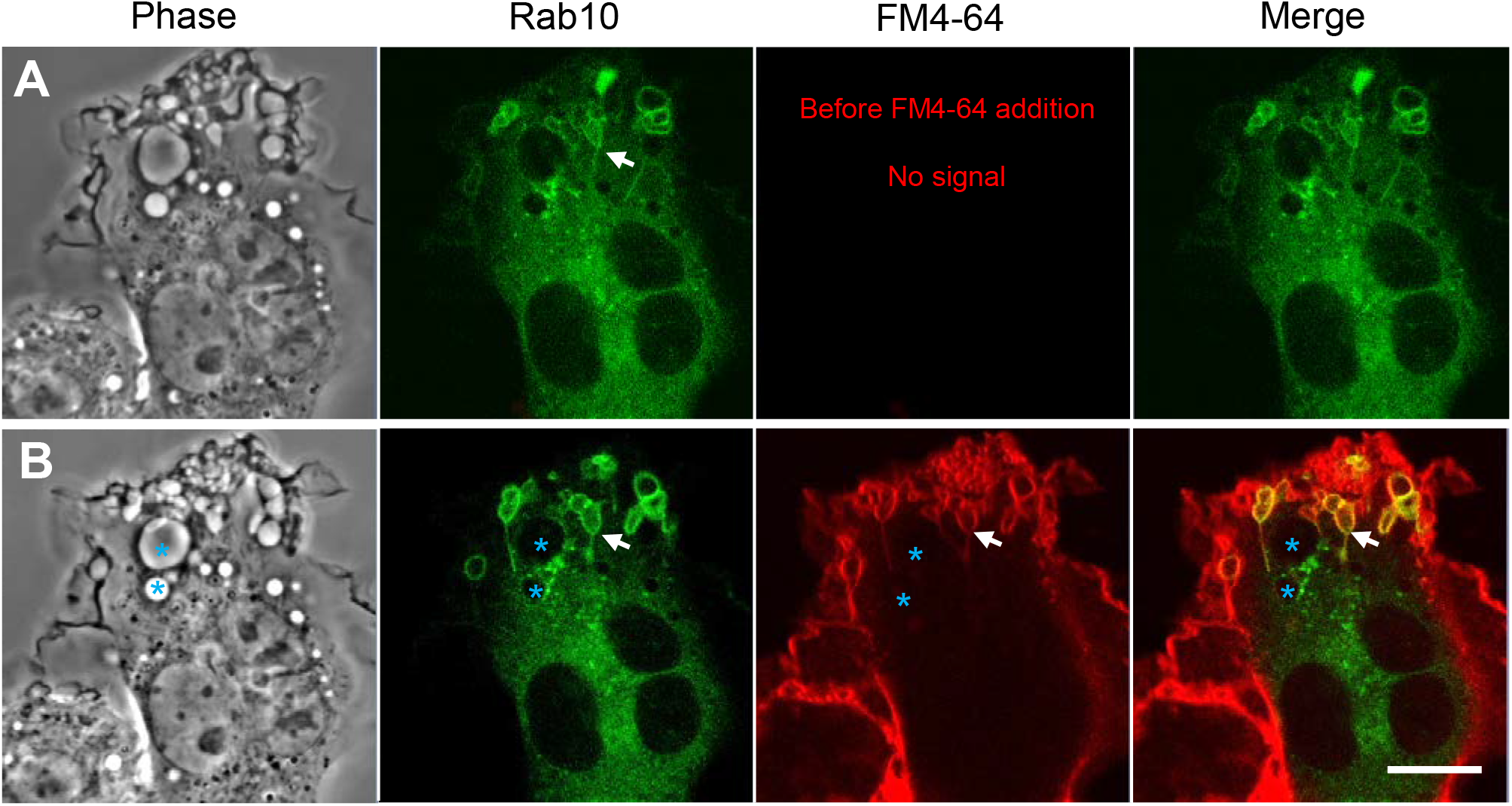
Rab10-positive macropinosome-like structures are unclosed macropinocytic cups (premacropinosomes). Rab10-positive macropinosome-like structures were induced by PA-Rac1 ON-OFF cycling. After confirming the formation of Rab10-positive macropinosome-like structures **(A)**, the FM4-64 membrane-impermeable dye was added to the medium to label the cell surface plasma membrane. The EGFP-Rab10 and FM4-64 images were acquired immediately after the FM4-64 addition **(B)**. Although phase-bright Rab10-negative macropinosomes (blue asterisks) were unlabeled with the FM4-64, Rab10-positive macropinocytic structures with extending tubules (arrow) were labeled with the FM4-64. Scale bar = 10 μm.

### Rab10-positive tubules that extend from premacropinosomes centripetally move along microtubules

Next, we examined whether Rab10-positive tubule movement was dependent on microtubules using RAW264 cells co-expressing EGFP-tubulin, mCherry-Rab10, and PA-Rac1. When Rab10-tubule formation was induced by PA-Rac1 ON-OFF cycling, long Rab10-positive tubules were observed to move along microtubules centripetally (Figures 4A, B. Supplementary Movies 4, 5). The long-distance vectorial movement of Rab10-positive tubules toward the perinuclear region was clearly disturbed when microtubules were disrupted by the addition of 3.3 μM nocodazole, a microtubule depolymerizer (Figure 4C. Supplementary Movie 6). However, the budding of short Rab10-positive tubules from premacropinosomes that displayed short-distance, non-vectorial movements was occasionally observed, even in the presence of nocodazole. These findings indicated that the vectorial movement of Rab10-positive tubules was highly dependent on microtubules, although the budding of Rab10-positive tubules from premacropinosomes may not require microtubules.

**Figure 4.**
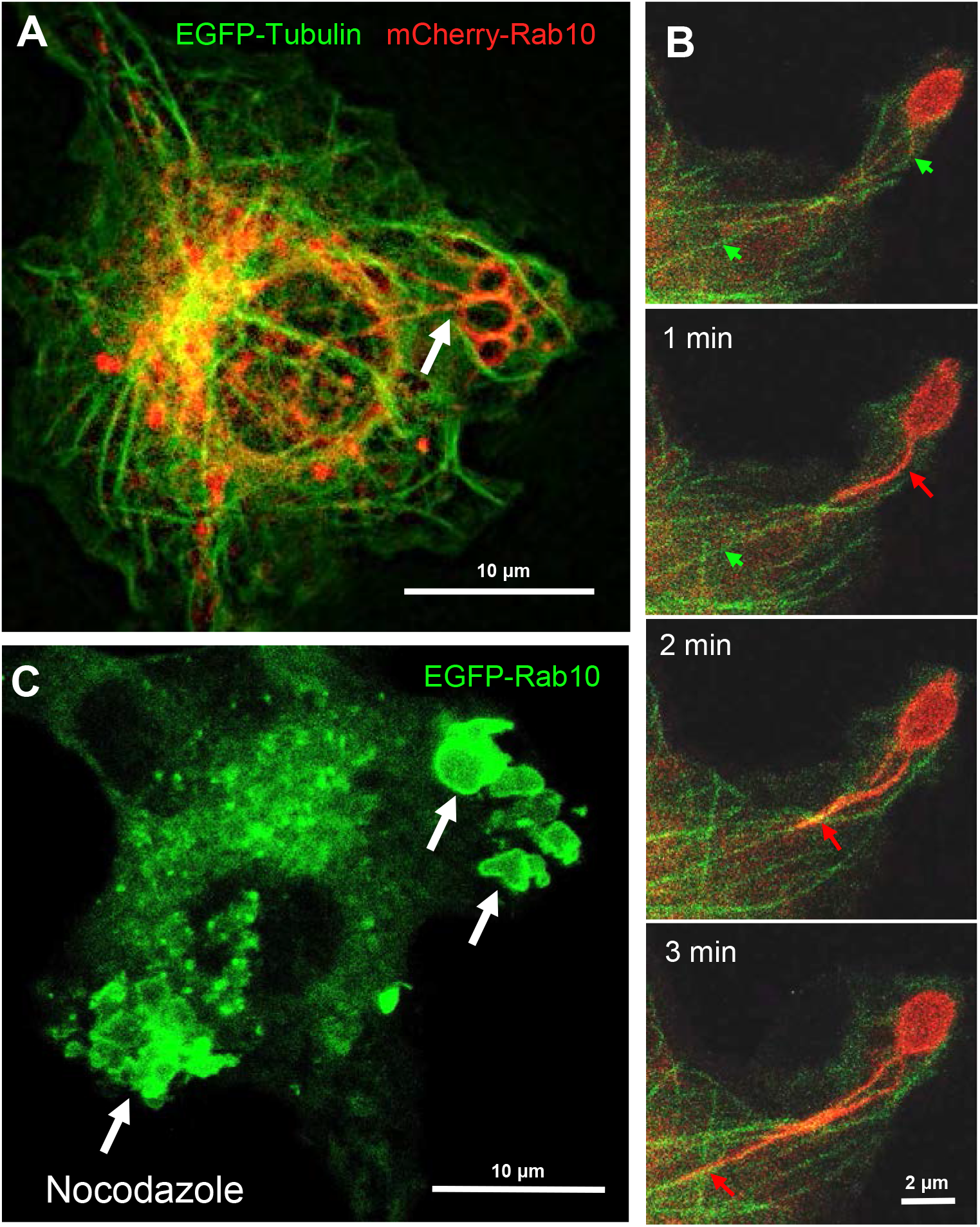
The retrograde movement of Rab10-positive tubules is dependent on microtubules. **(A)** Epifluorescence microscopy of mCherry-Rab10 and EGFP-tubulin in live RAW 264 cells during PA-Rac1 ON-OFF cycling. Arrow indicates a Rab10-positive tubule extended from a peripheral premacropinosome. Scale bar=10 μm. **(B)** Higher-magnification view of extending Rab10-positive tubules (red arrows) along microtubules (green arrows). Scale bar = 2 μm. **(C)** The extension of EGFP-Rab10 tubules was inhibited in nocodazole-treated RAW264 cells, while many Rab10-positive premacropinosomes (arrows) were formed. Scale bar = 10 μm. The corresponding movies are available in the Supplementary Materials (Movies 4–6).

### The Rab10 effectors EHBP1 and MICAL-L1 link Rab10 to EHD1 for membrane tubulation

Eps15 homology domain-binding protein 1 (EHBP1) is known as a Rab10 effector in the endocytic recycling pathway in *Caenorhabditis elegans*. It is reported that EHBP1 links EH domain-containing protein 1 (EHD1) to Rab10 to promote endosome tubulation (19,20). Therefore, we investigated the involvement of EHBP1 in Rab10-positive tubule formation from premacropinosomes in RAW264 cells co-expressing EGFP-EHBP1 and mCherry-Rab10. As expected, we observed the colocalization of Rab10 and EHBP1 in the premacropinosome and tubular structures induced by the PA-Rac1 ON-OFF photo-manipulation (Figure 5A, Supplementary Movie 7). EHD1 has membrane tubulation activity in an ATP-dependent manner and plays a role in the regulation of tubular recycling endosome trafficking to the plasma membrane (21). Consistently, EGFP-EHD1 was found to localize to the tubules budding from Rab10-positive premacropinosomes (Figure 5B, Supplementary Movie 8). Similarly, MICAL-L1, a Rab10 effector that links Rab10 to EHD1(22,23), was observed in Rab10-positive tubules (data not shown). These results suggested that EHPB1 and MICAL-L1 link Rab10 to EHD1 which may cause the tubulation of membranes.

**Figure 5.**
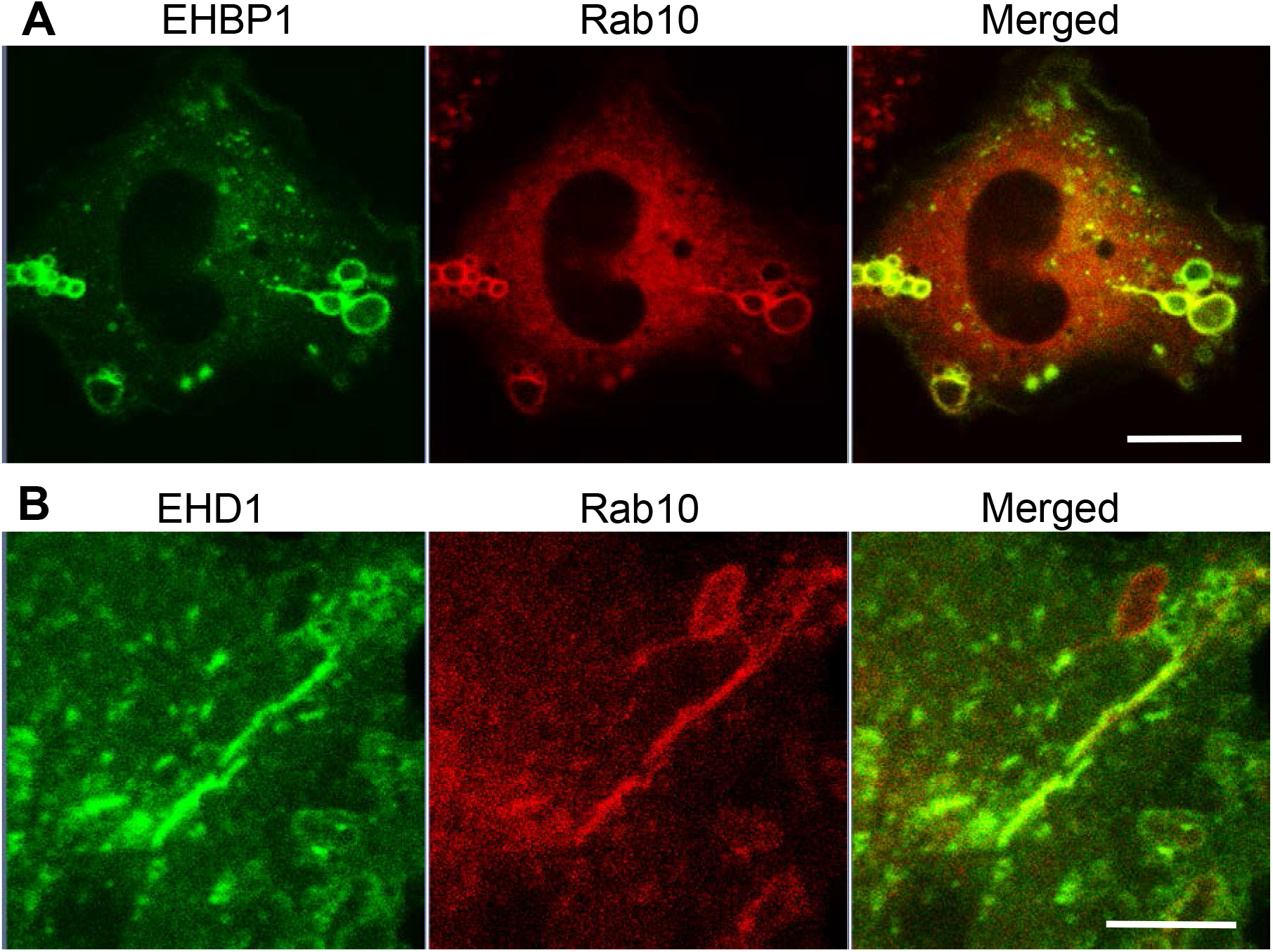
Localizations of EHBP1 and EHD1 on Rab10-positive premacropinosomes and tubules. **(A)** EHPB1 colocalized with Rab10 in premacropinosomes and tubules. Scale bar = 10 μm. **(B)** EHD1 predominantly localized on Rab10-positive tubules extended from premacropinosomes. Scale bar = 5 μm. Corresponding movies are available in the Supplementary Materials (Movies 7, 8).

### Rab10-positive tubular structures represent a novel endocytic pathway of membrane trafficking

Classical macropinosomes undergo a maturation process through the acquisition of Rab 5 and Rab7 before merging with lysosomes, where their contents are degraded. Therefore, we examined whether Rab10-positive tubules matured or fused with other endocytic compartments by performing a time-lapse image analysis in RAW264 cells co-expressing mCherry-Rab10 and various fluorescent protein-tagged endocytic marker proteins. Rab10-negative or faintly-positive compartments transiently acquire Rab5, an early endocytic protein. In contrast, Rab10-positive profiles do not become Rab5-positive (Figures 6A, B, Supplementary Movie 9). Also, Rab10-positive profiles were never observed to fuse with Rab7-positive late endocytic compartments or with LAMP1-positive lysosomal compartments. (Figures 7A, B, Supplementary Movies 10, 11). These findings indicated that these Rab10-positive profiles represented distinct entities from conventional endocytic pathways for lysosomal degradation.

**Figure 6.**
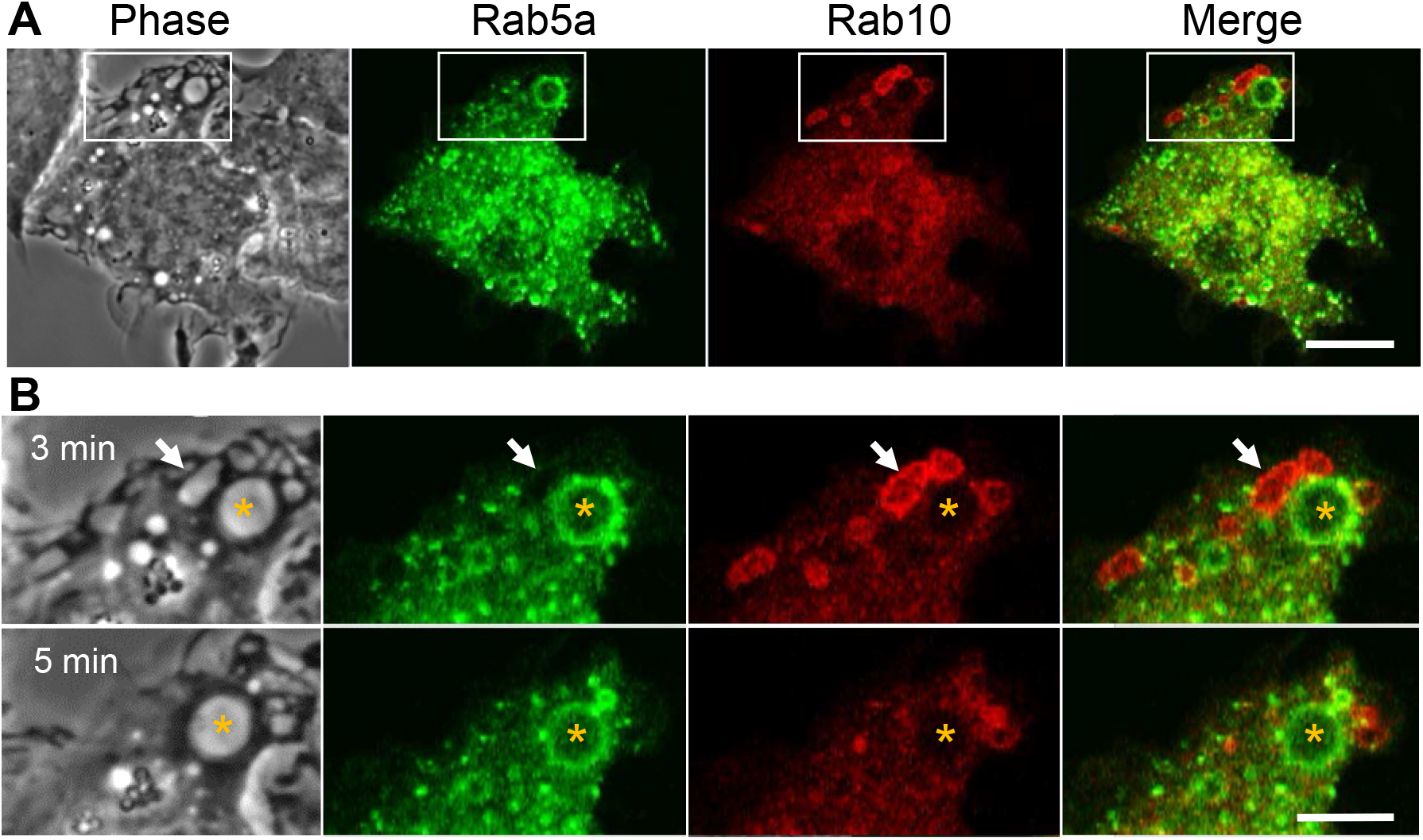
Rab5, an early endocytic marker, is not recruited to Rab10-positive premacropinosomes and tubules. **(A)** Live-cell microscopy of RAW264 cells expressing EGFP-Rab5a and mCherry-Rab10. Scale bar = 10 μm. **(B)** Enlarged micrographs of the boxed area in A. Selective frames at two-time points as indicated. Rab5a-positive macropinosomes are Rab10-negative (asterisk). Rab10-positive compartments disappear within a few minutes without becoming Rab5a-positive, suggesting that Rab10-positive compartments are distinct from those of the classical macropinocytic pathway. Scale bar = 5 μm. The corresponding movie is available in the Supplementary Materials (Movie 9).

**Figure 7.**
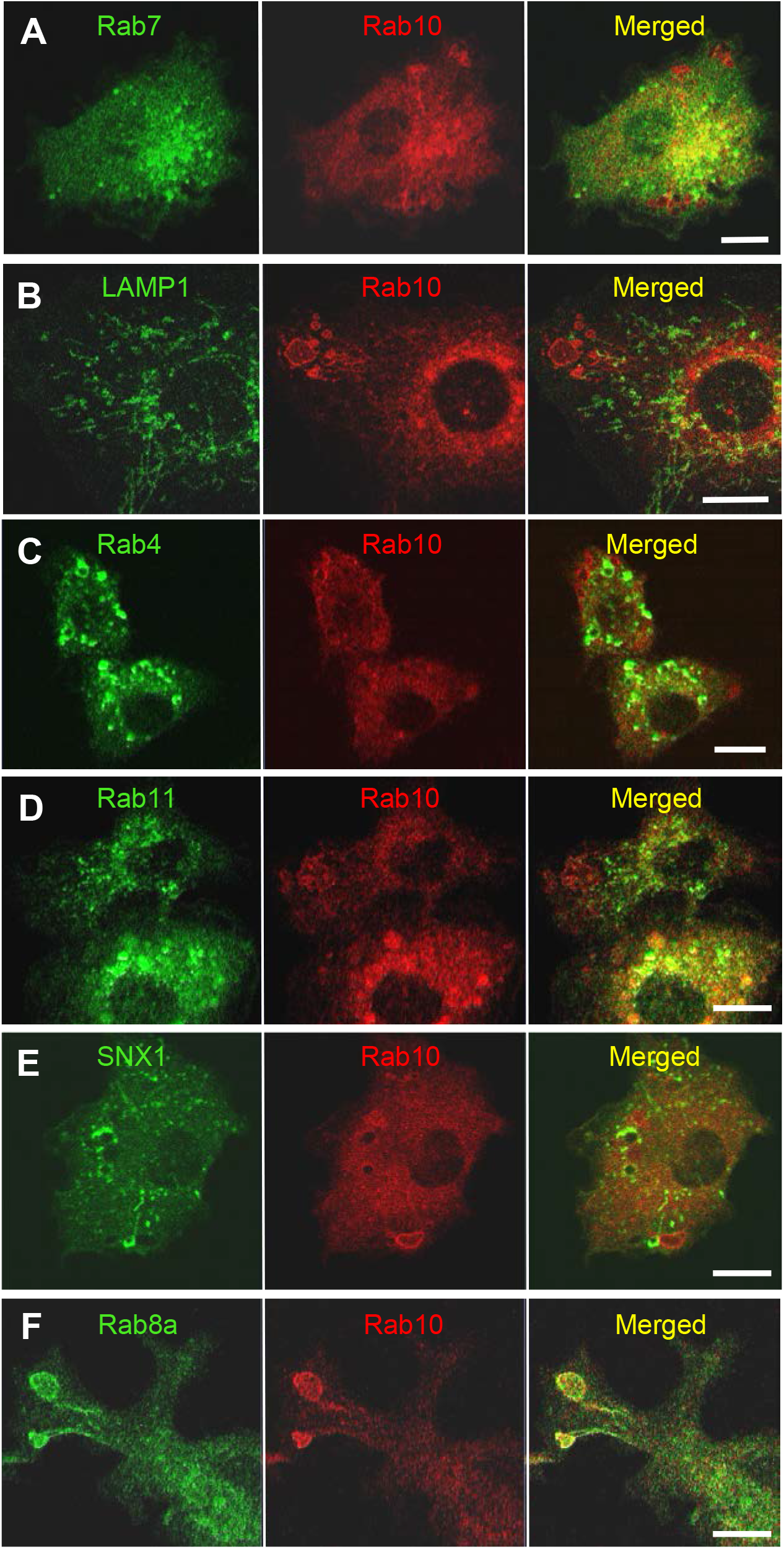
Rab10-positive premacropinosomes and tubules are distinct from conventional degradative endocytic or recycling pathways. Rab10-positive premacropinosomes and tubules are not labeled with the late endosome/lysosome markers, Rab7-**(A)** or LAMP1 **(B)**. The recycling endosomal markers, Rab4 **(C)**, Rab11 **(D)**, and sorting nexin 1 (SNX1) **(E)** are faintly observed in Rab10-positive premacropinosomes and tubules, although they are only transiently recruited to the Rab10-negative conventional macropinosomes. Only Rab8a colocalized with Rab10 in the premacropinosomes and extending tubules **(F)**. All scale bars = 10 μm. The corresponding movies are available in the Supplementary Materials (Movies 10–15).

Next, we examined the relationship between Rab10-positive tubules and recycling compartments using Rab4, which mediates fast recycling, or Rab11 which mediates slow recycling (24,25). Although EGFP-Rab11was slightly observed on the membranes of Rab10-positive premacropinosomes, Rab11 was not localized on Rab10-positive tubules. EGFP-Rab4 was also not observed on Rab10-positive premacropinosomes or tubules. These findings indicated that the Rab10-positive tubules that extended from premacropinosomes were not recycling compartments (Figures 7C, D, Supplementary Movies 12, 13). Sorting nexin 1 (SNX1), which contains BAR (Bin/Amphiphysin/Rvs) domains, localizes to early macropinosomes and tubules that extended from the early macropinosomes (26–28). However, we did not observe SNX1 localization on Rab10-positive compartments (Figure 7E, Supplementary Movie 14). Thus, Rab10-positive profiles likely exist as unique, non-degradative compartments that are distinct from canonical endosomes or recycling compartments. Among various Rab proteins that we examined, Rab8a, a Rab protein very closely related to Rab10, is found to be colocalized with Rab10 in the premacropinosomes and tubules (Figure 7F, Supplementary Movie 15).

### Rab10-positive premacropinosome and tubule formation is not dependent on PI3K

It is well-known that PI(3,4,5)P_3_ production by class I PI3K is critical for macropinosome closure during the process of classical macropinocytosis (29,30). However, the dependency of Rab10-positive premacropinosome and tubule formation on PI3K has not yet been determined. Therefore, we examined the PI3K-dependency of the Rab10-positive endocytic pathway using pharmacological inhibitors and mCitrine-fused Akt-PH domain (Akt-PH), which binds to PI(3,4,5)P_3_/PI(3,4)P_2_. In RAW 264 cells during PA-Rac1 ON-OFF cycling, the localization of Rab10 to macropinocytic cups or nascent macropinosomes was frequently observed. However, in most cases, the recruitment of Rab10 to these structures was transient and small in amount (Figure 8A, Supplementary Movies 16). In contrast, when the cells were treated with a PI3K inhibitor (10 μM LY294002 or 100 nM wortmannin), the remarkable accumulation of Rab10 in macropinocytic cups was frequently observed, although Rab10-negative macropinosomes were hardly seen (Figure 8B, Supplementary Movies 17). As a consequence, large macropinosomes rarely formed, whereas Rab10-positive tubule formation from premacropinosomes was not perturbed. Although Rab10 accumulation on the Akt-PH-positive macropinosomal membrane was not seen in RAW cells during PA-Rac1 ON-OFF cycling, Rab10-positive premacropinosomes without Akt-PH signals were frequently observed after treatment with PI3K inhibitors (Figures 8A, B Supplementary Movies 16, 17). These results suggested that Rab10-recruitment and Rab10-positive premacropinosome formation were PI3K-independent.

**Figure 8.**
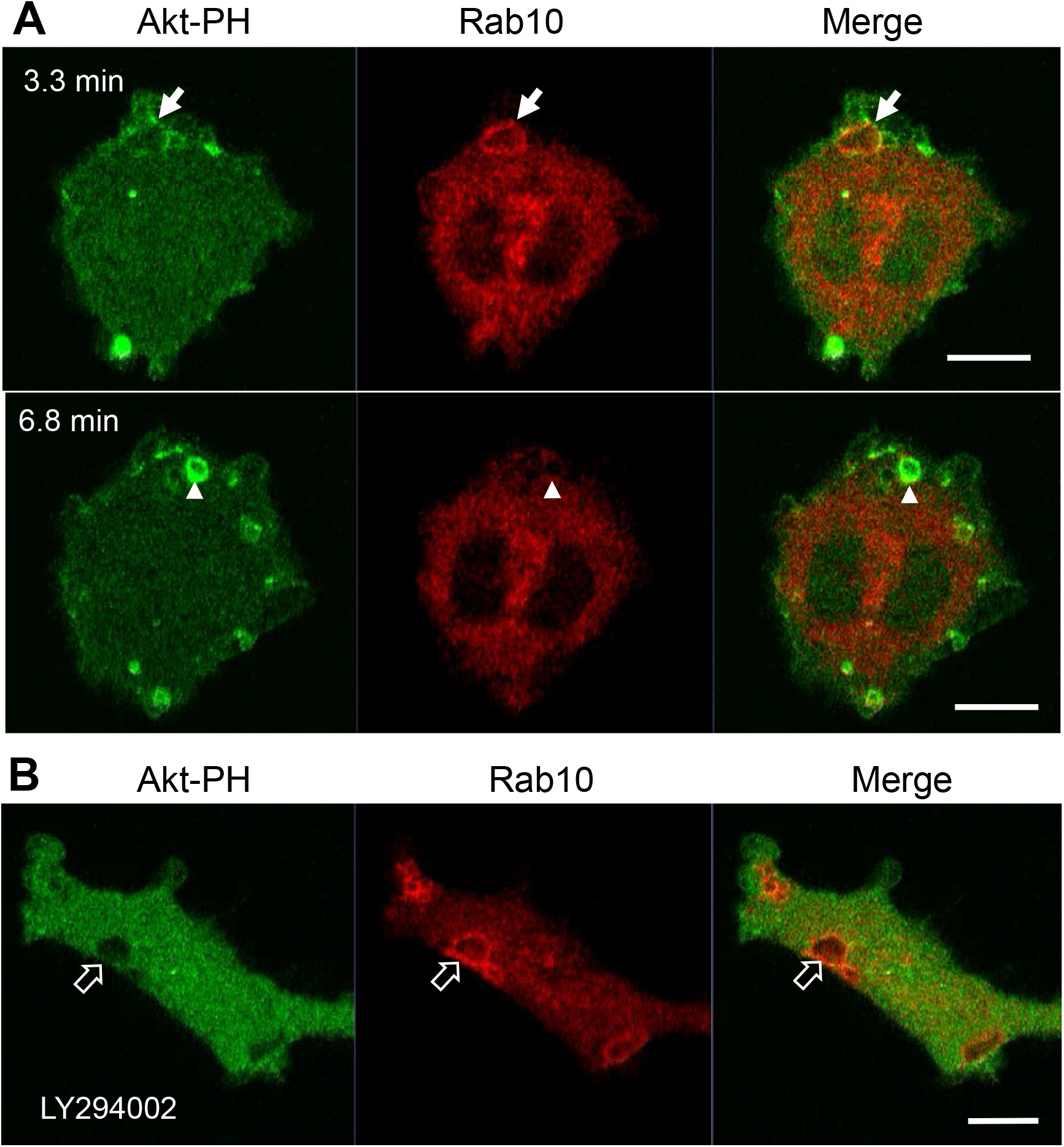
Premacropinosome formation and the recruitment of Rab10 are not PI3K-dependent. The PI(3,4,5)P_3_/PI(3,4)P_2_ production by class I PI3K was monitored through the expression of mCitrine-Akt-PH domain (Akt-PH, green) in a RAW264 cells during PA-Rac1 ON-OFF cycling **(A)** Although only small amounts of Akt-PH were observed on Rab10-positive premacropinosomes (arrows), Rab10-negative premacropinosomes were strongly positive for Akt-PH (arrowheads) at the timing of macropinosome closure. Elapsed time is shown in the frame. **(B)** In the presence of 10 μM LY294002, a PI3K inhibitor, the formation of Rab10-positive premacropinosomes was frequently seen (open arrows), whereas Akt-PH recruitment to the membrane was completely abolished. All scale bars = 10 μm. The corresponding movies are available in the Supplementary Materials (Movies 16, 17).

## Discussion

In this study, we first identified a novel endocytic pathway that diverges from canonical macropinocytosis using the optogenetic photo-manipulation of Rac1 activity. As we have reported previously (9), local PA-Rac1 activation induced the formation of premacropinosomes, which opened to the cell surface. Following the deactivation PA-Rac1, some of the premacropinosomes closed into intracellular macropinosomes, whereas the majority of these structures (~70%) disappeared within several minutes. Through the co-expression of PA-Rac1 and EGFP-Rab10, we found that Rab10-negative premacropinosomes became intracellular macropinosomes, whereas Rab10-enriched premacropinosomes shrank and disappeared without forming intracellular macropinosomes. Instead, Rab10-positive premacropinosomes transformed into Rab10-positive tubules that extended toward the perinuclear region. Because the premacropinosomes were unclosed, the fluid contents of the premacropinosomes were likely to be returned back to the extracellular fluid, and premacropinosome membranes can be efficiently transported because the Rab10-tubule features a much smaller diameter with a larger surface area/volume ratio than premacropinosome. Thus, the abundant and sustained localization of Rab10 on premacropinosomes may change the mode of endocytosis from fluid-phase to membrane flow.

Previous studies have shown that Rab10 is involved in membrane recycling to the plasma membrane, associated with tubular recycling endosomes in other cell types (15–17,20). Although the morphologies of these Rab10-positive tubular recycling endosomes are very similar to those observed in this study, the direction of movement for these tubular structures was the opposite of those observed here. Etoh and Fukuda revealed that Rab10-positive tubules extended from early endosomes in a microtubule-dependent manner and were anterogradely transported using the kinesin motors KIF13A and KIF13B (17). In contrast, the Rab10-positive tubules that extended from premacropinosomes were retrogradely transported along microtubules. Furthermore, the Rab10-positive tubules observed in this study were negative for Rab4 or Rab11, which are involved in fast recycling and slow recycling pathways, respectively (31), whereas the Rab10/Rab8a-positive tubules previously reported in other studies were associated with Rab4 or Rab11 proteins (13,17,32,33). Thus, the identified Rab10-positive tubules in this study are likely distinct from recycling tubules that traffic toward the plasma membrane. Curiously, no other early or late endocytic marker proteins, such as Rab5, Rab7, or LAMP1, were recruited to the Rab10-positive tubules observed in this study, which suggested that the Rab10-positive tubules that originate from premacropinosomes represent a non-degradative pathway that diverges from macropinocytosis.

Although SNX1, a tubulation protein having a BAR domain, is known to be localized on tubular recycling endosomes that derive from early macropinosomes (26–28), we could not detect SNX1 on the Rab10-positive tubules. Therefore, Rab10-positive tubules from premacropinosomes appear to be distinct from SNX1-positive recycling tubules that derive from early macropinosomes. Taken together, these findings suggested that the Rab10-positive tubules that extend from premacropinosomes toward the perinuclear Golgi region represent a previously undefined endocytic pathway.

Among the various Rab proteins that we examined, Rab8a showed a very similar localization pattern to Rab10 in the premacropinosomes and tubules. Because Rab8a/b are highly homologous to Rab10, they share common effectors, such as MICAL-L1 and EHBP1, in addition to upstream regulators, GEF, and GAP proteins. Therefore, Rab10 and Rab8 may be functionally redundant. Wall et al. (2017, 2019) reported that Rab8a is localized to macropinosomes and tubular compartments that elongate from macropinosomes in lipopolysaccharide (LPS)-stimulated RAW264.7 macrophages (34,35). Although they have regarded these Rab8-positive tubules as components of the conventional macropinocytosis pathway, the Rab8a-positive tubules observed in their studies may be identical to the Rab10-positive tubules observed in this study. In HeLaM cells, the knockout of Rab10 completely abolished Rab8- and MICAL-L1-positive tubular structures, whereas the Rab8a/b double-knockout did not perturb Rab10- and MICAL-L1-positive tubules (17). This study suggested that Rab10 plays a primary role in the formation of the tubular recycling endosome structure. Whether Rab10 and Rab8 play differential roles in macrophages should be clarified in future studies.

When exploring the underlying molecular mechanism responsible for Rab10-tubule formation, we found that EHBP1 and MICAL-L1, which are Rab10 effectors, both localized to Rab10-positive premacropinosomes and tubules. EHD1, a membrane tubulation protein, was also predominantly observed on Rab10-positive tubules. EHBP1 is known to link the EH domain-containing protein 1 (EHD1) to Rab10 to promote endosome tubulation (19,20). MICAL-L1 also links EHD1 to tubular recycling endosomes (22,23). Thus, Rab10 may recruit EHD1 to premacropinosomes through EHBP1 and/or MICAL-L1, causing membrane tubule budding from premacropinosomes.

Macropinosome formation is well-known to depend on PI(3,4,5)P_3_ production by class I PI3K (7,8,30,36). Live-cell imaging of the fluorescent protein-tagged Akt-PH domain demonstrated that PI(3,4,5)P_3_ levels in macropinocytic cup membranes rapidly increased at the timing of macropinosome closure (7,36). However, PI(3,4,5)P_3_ production was not observed upon the formation of Rab10-positive premacropinosomes. Furthermore, PI3K inhibition using pharmacological inhibitors did not reduce the formation of Rab10-positive premacropinosomes, whereas Rab10-negative macropinosome formation was drastically abolished. All Rab10-positive premacropinosomes were transformed into Rab10-positive tubules. These results suggested that the PI3K inhibition facilitates Rab10 localization to macropinocytic cups, leading to the transformation of premacropinosomes into tubules instead of macropinosome formation.

Recently, Liu et al. (2020) reported the transitional association of Rab10 with early macropinosomes in RAW macrophages. In their study, the LRRK2-mediated phosphorylation of Rab10 was shown to play a regulatory role in macropinosome maturation (18). However, they did not mention the formation of Rab10-positive tubules derived from the premacropinosomes that is persistently Rab10-positive. This inconsistency may be due to differences in the experimental conditions. Under the condition of PA-Rac1 photo-manipulation, the majority of the induced premacropinosomes transitioned into the Rab10-positive tubular endosome pathway, whereas LPS-induced macropinosomes underwent maturation in the classical macropinocytosis pathway. This difference is likely the reason why this Rab10-positive endocytic pathway has been overlooked or neglected. The identification of this novel pathway was a fortunate accident due to the use of innovative optogenetic technology. Importantly, we revealed that the Rab10-positive tubular endocytic pathway appears in macrophages even under near-physiological conditions, although the frequency is reduced under near-physiological conditions compared with PA-Rac1 photo-manipulation conditions. This finding suggested that the Rab10-positive endocytic pathway likely play a significant role under physiological and pathological conditions.

Although the function of Rab10 in macrophages remains obscure, Rab10 has recently been reported to be involved in the transport of Toll-like receptor 4 (TLR4) to the plasma membrane in macrophages (16). The Rab10-positive endocytic route identified in this study may be responsible for the membrane trafficking of TLR4 and other membrane proteins. Rab10 is thought to be involved primarily in transport from the Golgi complex to the plasma membrane, and our study has demonstrated, for the first time that Rab10 mediates a tubular membrane trafficking pathway that diverges from the canonical fluid-phase uptake by macropinocytosis.

Although the physiological, functional role of the Rab10-positive endocytic pathway remains unknown, the discovery of this unique non-degradative, endocytic pathway is likely to provide novel insights into the concepts of intracellular membrane trafficking. Intracellular pathways that avoids the lysosomal degradation system might be related to the pathophysiology of some metabolic diseases, and be utilized as a virus infection route. Further studies remain necessary to understand the functional role and molecular mechanisms underlying this novel endocytic pathway.

## Materials and Methods

### Chemicals and Plasmids

pEGFP-C1, pECFP-C1, and pmCherry-C1 were obtained from Clontech. pEGFP-Rab8a, pEGFP-Rab10, pmCherry-Rab10, pEGFP-Rab35, EGFP-tubulin, pEGFP-EHBP1, pEGFP-EHD1, pEGFP-MICAL-L1, and pEGFP-SNX1 were generated by amplifying the full-length open reading frames of each gene by polymerase chain reaction (PCR), followed by the cloning of the resultant PCR products into pEGFP-C1, or pmCherry-C1. The pmCitrine-Rab4, pmCitrine-Rab5, pmCitrine-Rab7, pmCitrine-LAMP1, and pmCitrine-Akt-PH domain were provided by Dr. Joel A. Swanson (University of Michigan). pEGFP-Rab11 was provided by Dr. Marino Zerial (Max Planck Institute). pTriEx/mCherry-PA-Rac1 was obtained from Dr. Klaus Hahn through Addgene (Plasmid #22027, Cambridge, MA). pECFP-PA-Rac1 was created by the insertion of the PA-Rac1 fragment into pECFP-C1.

Other reagents were purchased from Sigma-Aldrich (St. Louis, MO) or Nakalai Tesque (Kyoto, Japan) unless otherwise indicated.

### Cell culture and transfection

RAW264 cells were obtained from Riken Cell Bank (Tsukuba, Japan) and maintained in Dulbecco’s modified Eagle medium supplemented with 10% heat-inactivated fetal bovine serum and antibiotics (100 U/mL of penicillin and 0.1 mg/mL streptomycin) at 37°C in a humidified atmosphere containing 5% CO_2_. Cells were transfected with vectors using the Neon Transfection System (Life Technologies), according to the manufacturer’s protocol. Briefly, 100 μL of a RAW264 cell suspension (1.0 × 10^7^ cells/mL) in Buffer R was mixed with 1.0-3.0 mg of the indicated plasmids and electroporated once at 1,680 V for 20 ms. The cells were then seeded onto 25-mm circular coverslips in culture dishes containing the culture medium. At 10–24 h after transfection, the cells were subjected to live-cell imaging.

### Live-cell imaging and optogenetic photo-manipulation

RAW264 cells transfected with plasmids were cultured on 25-mm circular coverslips. The coverslip was assembled in an Attofluor cell chamber (A7816, Molecular Probes, Eugene, OR) filled with Ringer’s buffer (RB), consisting of 155 mM NaCl, 5 mM KCl, 1 mM MgCl_2_, 2 mM Na_2_HPO_4_, 10 mM glucose, 10 mM HEPES and 0.5 mg/mL bovine serum albumin (BSA) at pH 7.2. The chamber was settled onto a thermo-controlled stage (Tokai Hit, Shizuoka, Japan) attached to a confocal laser microscope (Zeiss LSM700) or an epifluorescence microscope (Leica DMI6000B).

The optogenetic photo-manipulation of PA-Rac1 activity was performed as previously described (9,37). Briefly, cells were transfected with pTriEx/mCherry-or ECYP-PA-Rac1 (38) and observed under the LSM700 confocal microscope controlled by ZEN (Zeiss) By illuminating a blue-light laser (445 or 488 nm wavelength) to the cells expressing PA-Rac1 under the indicated conditions, PA-Rac1 was activated through the conformational change of the light oxygen voltage 2 domain (LOV2) of PA-Rac1 in either a local area or the whole-cell region. PA-Rac1 can be reversibly deactivated by turning off the blue-light illumination (38). To observe the dynamics of a red fluorescent protein-tagged protein during PA-Rac1 photo-manipulation, ECYP-PA-Rac1 was employed. In some optogenetic photo-manipulation experiments were performed using the Leica DMI6000B automated epifluorescence microscopy system, controlled by MetaMorph software (Molecular Devices) (37). The acquired fluorescence images were processed using the Safir denoising software (INRIA) and the nearest neighbor deconvolution algorism of the MetaMorph software. Microscopic images are representative of ≥7 cells from at least three separate experiments.

### Data presentation and statistical analysis

Quantitation of macropinosome and premacropinosome formation was performed by counting phase-bright macropinosomes and Rab10-positive premacropinosomes larger than 2 μm in diameter using time-lapse microscopic movies. The number of macropinosomes/premacropinosomes formed during 10 min/cell was calculated. Quantitative data are expresses as the means ± standard deviations (SD, n≥7 cells from at least three independent experiments). Significant differences in mean values were determined by a two-tailed unpaired *t-test. P*-values less than 0.05 were considered statistically significant.

## Author Contributions

Conceived and designed the experiments: KK, NA. Performed the experiments and collected image data: KK, AN, SM. Analyzed the data: KK, NA. Contributed reagents/materials/analysis tools: KK, AN, SM, YE. Wrote and edited the manuscript; NA, KK. All authors contributed to the article and approved the submitted version.

## Acknowledgments

We are grateful to Drs. Joel A. Swanson (University of Michigan), Marino Zerial (Max Planck Institute), and Klaus Hahn (University of North Carolina) for providing plasmids; and to Mr. Toshitaka Nakagawa, Mr. Kazuhiro Yokoi, and Ms. Yukiko Iwabu for their skillful assistance. We also wish to thank Yuri Manabe, Akira Ikawa, Yusei Nonomiya, Daisuke Katayama, and Masana Hayashi, for their contribution to this research project in the undergraduate medical science research program at the Araki laboratory.

---

**Supplementary Figure S1.**
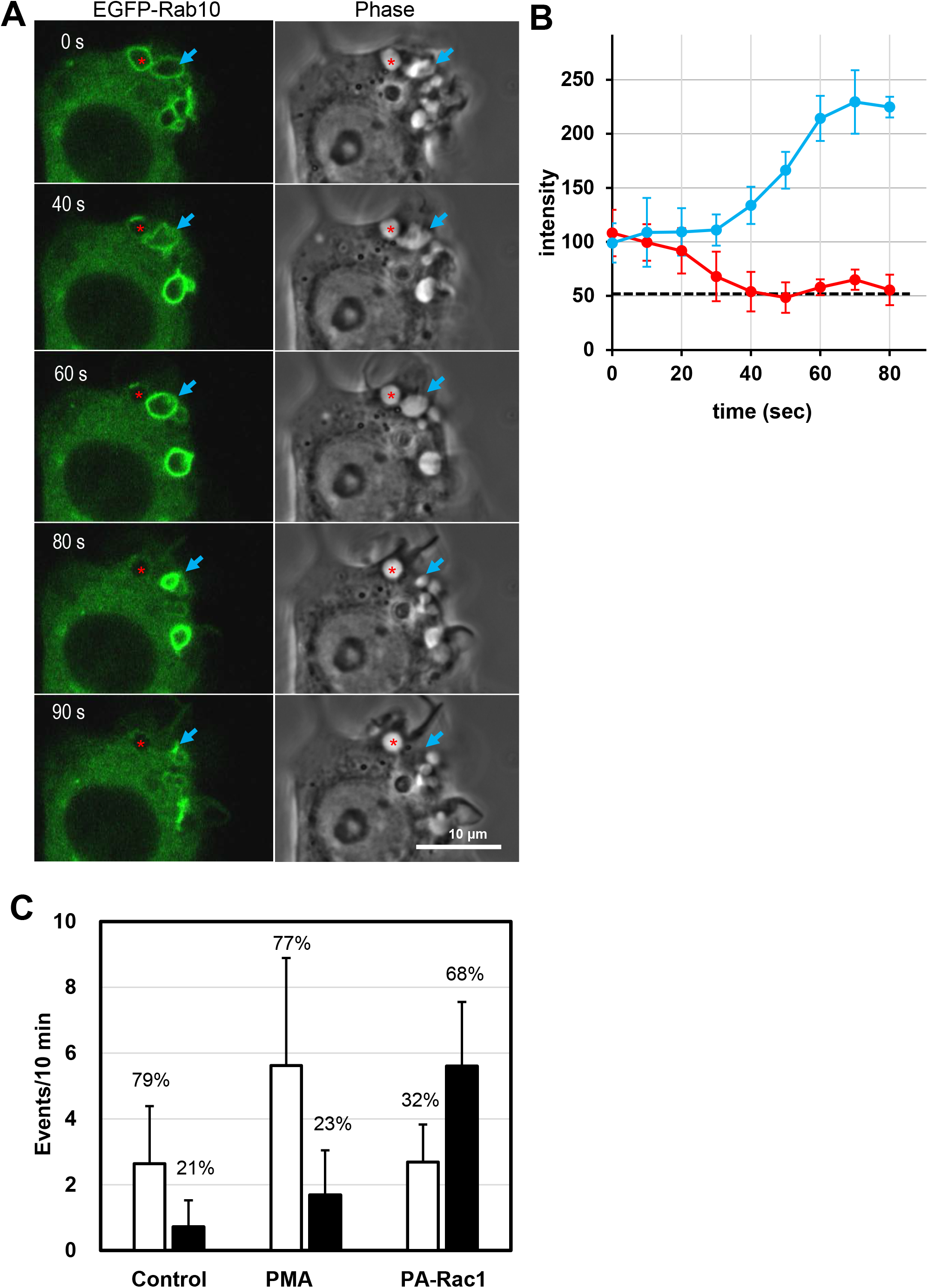
**(A)** Confocal live-cell imaging of EGFP-Rab10 in RAW264 cells during PA-Rac1 ON-OFF cycling. Selected images from a time-lapse movie of EGFP-Rab10 (left column) and phase-contrast (right column) images. Scale bar = 10 μm. **(B)** Graph showing the changes in the fluorescence intensity of EGFP-n intensity of EGFP-Rab10 was measured at four points on the macropinocytic membrane. Data are expressed as the mean ± SD (n=4 points). The red line indicates the classical macropinocytosis pathway (asterisks in **A**). The blue line indicates the Rab10-positive endocytic pathway (arrows in **A**). **(C)** The formation frequency of Rab10-negative macropinosomes (white bars) and Rab10-positive premacropinosomes (filled bars) was quantified by counting the events in time-lapse movies of EGFP-Rab10 and phase-contrast images of control RAW264 cells (Control), PMA-stimulated RAW264 cells (PMA), and cells stimulated with PA-Rac1 ON-OFF cycling (PA-Rac1). Data are expressed as means (events/10 min) ± SD (n≥6 cells). The percentages of each event in all macropinocytic events are shown above each bar.

## Legends of Movies

**Movie 1.** Optogenetic control of PA-Rac1 activity in RAW264 cells. The boxed area was locally irradiated with a blue laser to activate Rac1 (PA-Rac1 ON); then, the blue laser was turned off to deactivate Rac1 (PA-Rac1 OFF). ×80 speed. Scale bar = 10 μm.

**Movie 2.** Time-lapse movie showing the formation of Rab10-positive premacropinosomes and tubules in RAW264 macrophages expressing EGFP-Rab10 by PA-Rac1 ON-OFF cycling. ×80 speed. Scale bar = 10 μm.

**Movie 3.** EGFP-Rab10 and phase-contrast overlay images showing the formation of short-lifetime, EGFP-Rab10-positive premacropinosomes and Rab10-negative classical macropinosomes in a PMA-stimulated RAW264 cell. ×80 speed. Scale bar = 10 μm.

**Movie 4.** Time-lapse movie showing dynamics of mCherry-Rab10 and EGFP-tubulin in live RAW264 cells during PA-Rac1 ON-OFF cycling. ×30 speed. Scale bar = 10 μm.

**Movie 5.** High-magnification movie showing extending Rab10-positive tubules (red) along microtubules (green). × 80 speed, Scale bar = 2 μm.

**Movie 6.** Time-lapse movie showing dynamics of EGFP-Rab10 in a nocodazole-treated RAW264 cells during PA-Rac1 ON-OFF cycling. ×80 speed. Scale bar = 10 μm.

**Movie 7.** Dual-color time-lapse movie showing that EGFP-EHPB1 (green) is found to colocalize with mCherry-Rab10 (red) in premacropinosomes and tubules. ×80 speed. Scale bar = 10 μm.

**Movie 8.** Dual-color time-lapse movie showing that EGFP-EHD1 (green) is predominantly localized on mCherry-Rab10 (red)-positive tubules extending from premacropinosomes. ×80 speed. Scale bar = 5 μm.

**Movie 9.** Dual-color time-lapse movie showing that EGFP-Rab5a (green) and mCherry-Rab10 (red) localize to distinct compartments. Although the Rab10-negative, Rab5a-temporarily positive compartment remains a phase-bright macropinosome (arrow), mCherry-Rab10-positive compartments disappear within a few minutes without becoming Rab5a-positive. The merge image was overlaid with the corresponding phase-contrast image (right). ×80 speed. Scale bar = 10 μm.

**Movie 10.** Dual-color time-lapse movie showing that mCherry-Rab10 (red)-positive tubules extending from Rab10-positive premacropinosomes do not obtain mCitrine-Rab7 (green). ×80 speed. Scale bar = 10 μm.

**Movie 11.** Dual-color time-lapse movie showing that mCherry-Rab10 (red)-positive premacropinosomes and extending tubules do not merge with mCitrine-LAMP1 (green). ×80 speed. Scale bar=10 μm.

**Movie 12.** Dual-color time-lapse movie showing that mCherry-Rab10 (red)-positive premacropinosomes and extending tubules are distinct compartments from EGFP-Rab4 (green)-positive fast recycling endosomal compartments. ×80 speed. Scale bar = 10 μm.

**Movie 13.** Dual-color time-lapse movie showing that mCherry-Rab10 (red)-positive premacropinosomes and extending tubules are distinct compartments from EGFP-Rab11 (green)-positive recycling endosomal compartments. ×80 speed. Scale bar = 10 μm.

**Movie 14.** Dual-color time-lapse movie showing that mCherry-Rab10 (red)-positive premacropinosomes and extending tubules are distinct compartments from EGFP-SNX1 (green)-positive recycling tubules that extend from conventional macropinosomes. ×80 speed. Scale bar = 10 μm.

**Movie.15.** Dual-color time-lapse movie showing that EGFP-Rab8a (green) and mCherry-Rab10 colocalize on premacropinosomes and extending tubules. ×80 speed. Scale bar = 10 μm.

**Movie 16.** Dual-color time-lapse movie showing PI(3,4,5)P_3_/PI(3,4)P_2_ production, monitored by the mCitrine-Akt PH domain (green) in a RAW264 cell during PA-Rac1 ON-OFF cycling. Although only small amounts of Akt-PH are observed on Rab10-positive premacropinosomes (green arrow), Rab10-negative premacropinosomes are strongly positive for Akt-PH (red arrow). ×80 speed. Scale bar=10 μm.

**Movie 17.** Dual-color time-lapse movie showing that LY294002 does not inhibit Rab10-positive premacropinosome formation, while the recruitment of mCitrine-Akt-PH (green) to Rab10-positive premacropinosomal membranes was completely abolished. Arrows indicate the formation of Rab10-positive premacropinosomes by PA-Rac1 ON-OFF cycling. ×80 speed. Scale bar = 10 μm.

## References

1. Swanson JA, Watts C. Macropinocytosis. Trends Cell Biol (1995) 5:424–428. doi:10.1016/S0962-8924(00)89101-1

2. Kerr MC, Teasdale RD. Defining Macropinocytosis. Traffic (2009) 10:364–371. doi:10.1111/j.1600-0854.2009.00878.x

3. Sallusto F, Cella M, Danieli C, Lanzavecchia A. Dendritic cells use macropinocytosis and the mannose receptor to concentrate macromolecules in the major histocompatibility complex class II compartment: downregulation by cytokines and bacterial products. J Exp Med (1995) 182:389–400. doi:10.1084/jem.182.2.389

4. Commisso C, Davidson SM, Soydaner-Azeloglu RG, Parker SJ, Kamphorst JJ, Hackett S, Grabocka E, Nofal M, Drebin JA, Thompson CB, et al. Macropinocytosis of protein is an amino acid supply route in Ras-transformed cells. Nature (2013) 497:633–7. doi:10.1038/nature12138

5. Zeineddine R, Yerbury JJ. The role of macropinocytosis in the propagation of protein aggregation associated with neurodegenerative diseases. Front Physiol (2015) 6:277. doi:10.3389/fphys.2015.00277

6. Mercer J, Helenius A. Virus entry by macropinocytosis. Nat Cell Biol (2009) 11:510–20. doi:10.1038/ncb0509-510

7. Araki N, Egami Y, Watanabe Y, Hatae T. Phosphoinositide metabolism during membrane ruffling and macropinosome formation in EGF-stimulated A431 cells. Exp Cell Res (2007) 313:1496–507. doi:10.1016/j.yexcr.2007.02.012

8. Egami Y, Taguchi T, Maekawa M, Arai H, Araki N. Small GTPases and phosphoinositides in the regulatory mechanisms of macropinosome formation and maturation. Front Physiol (2014) 5:374. doi:10.3389/fphys.2014.00374

9. Fujii M, Kawai K, Egami Y, Araki N. Dissecting the roles of Rac1 activation and deactivation in macropinocytosis using microscopic photo-manipulation. Sci Rep (2013) 3:2385. doi:10.1038/srep02385

10. Stenmark H. Rab GTPases as coordinators of vesicle traffic. Nat Rev Mol Cell Biol (2009) 10:513–525. doi:10.1038/nrm2728

11. Pfeffer S, Aivazian D. Targeting Rab GTPases to distinct membrane compartments. Nat Rev Mol Cell Biol (2004) doi:10.1038/nrm1500

12. Babbey CM, Ahktar N, Wang E, Chen CCH, Grant BD, Dunn KW. Rab10 regulates membrane transport through early endosomes of polarized Madin-Darby Canine Kidney cells. Mol Biol Cell (2006) doi:10.1091/mbc.E05-08-0799

13. Chua CEL, Tang BL. Rab 10—a traffic controller in multiple cellular pathways and locations. J Cell Physiol (2018) doi:10.1002/jcp.26503

14. Sano H, Eguez L, Teruel MN, Fukuda M, Chuang TD, Chavez JA, Lienhard GE, McGraw TE. Rab10, a Target of the AS160 Rab GAP, Is Required for Insulin-Stimulated Translocation of GLUT4 to the Adipocyte Plasma Membrane. Cell Metab (2007) doi:10.1016/j.cmet.2007.03.001

15. Shi A, Liu O, Koenig S, Banerjee R, Chen CC-H, Eimer S, Grant BD. RAB-10-GTPase-mediated regulation of endosomal phosphatidylinositol-4,5-bisphosphate. Proc Natl Acad Sci U S A (2012) 109:E2306–15. doi:10.1073/pnas.1205278109

16. Wang D, Lou J, Ouyang C, Chen W, Liu Y, Liu X, Cao X, Wang J, Lu L. Ras-related protein Rab10 facilitates TLR4 signaling by promoting replenishment of TLR4 onto the plasma membrane. Proc Natl Acad Sci (2010) 107:13806–13811. doi:10.1073/pnas.1009428107

17. Etoh K, Fukuda M. Rab10 regulates tubular endosome formation through KIF13A and KIF13B motors. J Cell Sci (2019) doi:10.1242/jcs.226977

18. Liu Z, Xu E, Zhao HT, Cole T, West AB. LRRK2 and Rab10 coordinate macropinocytosis to mediate immunological responses in phagocytes. EMBO J (2020)e104862. doi:10.15252/embj.2020104862

19. Shi A, Chen CCH, Banerjee R, Glodowski D, Audhya A, Rongo C, Grant BD. EHBP-1 functions with RAB-10 during endocytic recycling in Caenorhabditis elegans. Mol Biol Cell (2010) doi:10.1091/mbc.E10-02-0149

20. Wang P, Liu H, Wang Y, Liu O, Zhang J, Gleason A, Yang Z, Wang H, Shi A, Grant BD. RAB-10 promotes EHBP-1 bridging of filamentous actin and tubular recycling endosomes. PLOS Genet (2016) 12:e1006093. doi:10.1371/journal.pgen.1006093

21. Lee S, Uchida Y, Wang J, Matsudaira T, Nakagawa T, Kishimoto T, Mukai K, Inaba T, Kobayashi T, Molday RS, et al. Transport through recycling endosomes requires EHD1 recruitment by a phosphatidylserine translocase. EMBO J (2015) 34:669–88. doi:10.15252/embj.201489703

22. Sharma M, Srinivas Panapakkam Giridharan S, Rahajeng J, Caplan S, Naslavsky N. MICAL-L1. Commun Integr Biol (2010) 3:181–183. doi:10.4161/cib.3.2.10845

23. Sharma M, Giridharan SSP, Rahajeng J, Naslavsky N, Caplan S. MICAL-L1 links EHD1 to tubular recycling endosomes and regulates receptor recycling. Mol Biol Cell (2009) 20:5181–5194. doi:10.1091/mbc.E09-06-0535

24. Pfeffer S. Membrane domains in the secretory and endocytic pathways. Cell (2003) doi:10.1016/S0092-8674(03)00118-1

25. Zerial M, McBride H. Rab proteins as membrane organizers. Nat Rev Mol Cell Biol (2001) 2:107–117. doi:10.1038/35052055

26. Buckley CM, King JS. Drinking problems: mechanisms of macropinosome formation and maturation. FEBS J (2017) doi:10.1111/febs.14115

27. Kerr MC, Lindsay MR, Luetterforst R, Hamilton N, Simpson F, Parton RG, Gleeson PA, Teasdale RD. Visualisation of macropinosome maturation by the recruitment of sorting nexins. J Cell Sci (2006) 119:3967–3980. doi:10.1242/jcs.03167

28. Lim JP, Gleeson PA, Teasdale RD. SNX5 is essential for efficient macropinocytosis and antigen processing in primary macrophages. Biol Open (2012) 1:904–914. doi:10.1242/bio.20122204

29. Cox D, Tseng CC, Bjekic G, Greenberg S. A requirement for phosphatidylinositol 3-kinase in pseudopod extension. J Biol Chem (1999) 274:1240–7. Available at: http://www.ncbi.nlm.nih.gov/entrez/query.fcgi?cmd=Retrieve&db=PubMed&dopt=Citation&list_uids=9880492 [Accessed July 14, 2014]

30. Araki N, Johnson MT, Swanson JA. A role for phosphoinositide 3-kinase in the completion of macropinocytosis and phagocytosis by macrophages. J Cell Biol (1996) 135:1249–60. doi:10.1083/jcb.135.5.1249

31. Grant BD, Donaldson JG. Pathways and mechanisms of endocytic recycling. Nat Rev Mol Cell Biol (2009) doi:10.1038/nrm2755

32. Lucken-Ardjomande Häsler S, Vallis Y, Pasche M, McMahon HT. GRAF2, WDR44, and MICAL1 mediate Rab8/10/11–dependent export of E-cadherin, MMP14, and CFTR ΔF508. J Cell Biol (2020) 219: doi:10.1083/jcb.201811014

33. Chen Y, Wang Y, Zhang J, Deng Y, Jiang L, Song E, Wu XS, Hammer JA, Xu T, Lippincott-Schwartz J. Rab10 and myosin-Va mediate insulin-stimulated GLUT4 storage vesicle translocation in adipocytes. J Cell Biol (2012) 198:545–560. doi:10.1083/jcb.201111091

34. Wall AA, Luo L, Hung Y, Tong SJ, Condon ND, Blumenthal A, Sweet MJ, Stow JL. Small GTPase Rab8a-recruited phosphatidylinositol 3-kinase γ regulates signaling and cytokine outputs from endosomal Toll-like receptors. J Biol Chem (2017) 292:4411–4422. doi:10.1074/jbc.M116.766337

35. Wall AA, Condon ND, Luo L, Stow JL. Rab8a localisation and activation by Toll-like receptors on macrophage macropinosomes. Philos Trans R Soc B Biol Sci (2019) doi:10.1098/rstb.2018.0151

36. Yoshida S, Hoppe AD, Araki N, Swanson JA. Sequential signaling in plasma-membrane domains during macropinosome formation in macrophages. J Cell Sci (2009) 122:3250–61. doi:10.1242/jcs.053207

37. Araki N, Ikeda Y, Kato T, Kawai K, Egami Y, Miyake K, Tsurumaki N, Yamaguchi M. Development of an automated fluorescence microscopy system for photomanipulation of genetically encoded photoactivatable proteins (optogenetics) in live cells. Microscopy (2014) 63:255–260. doi:10.1093/jmicro/dfu003

38. Wu YI, Frey D, Lungu OI, Jaehrig A, Schlichting I, Kuhlman B, Hahn KM. A genetically encoded photoactivatable Rac controls the motility of living cells. Nature (2009) 461:104–8. doi:10.1038/nature08241

